# Repeated Decision Stumping Distils Simple Rules from Single Cell Data

**DOI:** 10.1101/2020.09.08.288662

**Authors:** Ivan A. Croydon Veleslavov, Michael P.H. Stumpf

## Abstract

Here we introduce repeated decision stumping, to distill simple models from single cell data. We develop decision trees of depth one – hence ‘stumps’ – to identify in an inductive manner, gene products involved in driving cell fate transitions, and in applications to published data we are able to discover the key-players involved in these processes in an unbiased manner without prior knowledge. The approach is computationally efficient, has remarkable predictive power, and yields robust and statistically stable predictors: the same set of candidates is generated by applying the algorithm to different subsamples of the data.

## Background

The development, testing, and iterative improvements of models and hypotheses are cornerstones of scientific practice. There is, however, an inevitable trade-off between predictive power and interpretability in any modelling of real-world data and processes[1]. This is especially true for deep neural networks or reinforcement learning which are capable of capturing complex and non-linear relationships among large training data. But understanding the basis of these predictions – or distilling mechanistic insights from them – has often proved hard[2]. While recent applications to single-cell data have demonstrated the predictive power of these methods[3, 4], applications of machine and statistical learning to this type of data are also revealing how difficult it is to gain concrete mechanistic biological insights from these black-box approaches.

A model with high predictive performance in assigning cell types, for example, may in some contexts be less useful or interesting than a model with worse predictive performance that instead offers some mechanistic insights as to why a certain cell type is realised[5, 1]. Even a small insight, a “rule of thumb” or engineering heuristic, of what drives a cell-fate transition[6, 7, 8, 9, 10] can provide useful and interpretable insights into the fundamental biological processes. Distilling such simple rules from complex data is our primary motivation here.

Single cell technologies have opened up new avenues for the analysis of cellular processes[11, 12, 13, 2]. Importantly, they have revealed widespread cell-to-cell variability of apparently phenotypically identical cells at the molecular level. Transcriptomic analyses account for the majority of single cell data, and single cell transcriptomics has become widely deployed in e.g. stem cell and developmental biology, but also in immunology and cancer research. Changes at the transcriptome level are widely seen as indicative of, and even causal for, cell fate decision making and progression of cells along the differentiation path[14, 15].

The riches contained in these data, do not reveal themselves easily, however, as a host of problems plague analysisandinterpretation[12, 2]: experimental(undesirable) noise interfereswiththemolecularnoise(which holds signatures of molecular processes); the high dimensionality of the data poses considerable statistical challenges. Models that learn to make robust predictions for cell fates under such noise carry clear motivation[16]. Further, if we are able to design these models to also provide basic biological insight, we will have a pragmatic tool, based on decision stumps[17, 18, 19], with which to (i) study high-dimensional developmental dynamics; or (ii) develop engineering strategies e.g. to control cellular behaviour [20, 21, 22].

Here we introduce a new algorithm, *Repeated Decision Stumping* (ReDX), that exploits the simple structure of decision tree models to distill highly interpretable but still predictive insights from single cell data: single genes or sets of genes that show particular statistical association with developmental events. This approach complements traditional studies that e.g. follow sets of genes in a hypothesis driven manner; and data-driven approaches that approach transcriptomic data without a specific hypothesis. ReDX sits between these two types of algorithms and allows us to generate new interpretable hypotheses and mechanistic models in a data-driven framework.

## Results

In the following, we demonstrate the effectiveness and pragmatism of this approach in identifying testable hypotheses for a range of developmental contexts. We are particularly interested in finding genes with expression patterns that align well with changes in cell state. These transitions are, of course, initiated and orchestrated by gene regulatory interactions [23, 11, 12]; but identification of genes that show a particularly strong association with a cell fate decision making process or transition can help with improving our mechanistic understanding, and the development or improvement of models describing such processes. We show that the models learnt by ReDX exhibit high predictive performance on withheld test data; furthermore the simplicity of the model structures allows us to obtain interpretable insights into the complex developmental dynamics. We also devise a metric, *γ*, for quantifying the stability [24, 25] of this feature selection under random permutations of the training data, and use it to demonstrate the robustness of the resulting predictions.

### Identifying genes associated with mouse neuronal precursors

We first apply ReDX to the Stumpf *et al*. [26, 23] data on murine neural differentiation. Here mouse embryonic stem (ES) cells were profiled using RT-q-PCR along their commitment to neuronal precursor (NP) cells, via an epiblast-like intermediate cell stage (EPI). For each successive developmental boundary captured in the data (ES→EPI; EPI→NP), we apply ReDX to identify the topmost informative genes for discriminating between the cell classes.

The results show that ReDX is able to learn simple models that characterise each transition well - in terms of predictive performance on the withheld test data - whilst incorporating genes already implicated in murine neuronal commitment [Table 1].

**Table 1:** Results of ReDX applied to murine neuronal cell fate commitment, with 5 iterations. The feature name and chosen threshold is presented, alongside each model’s performance on withheld data. The direction of the prediction is also provided, with respect to the established lineage ES→EPI→NPC.

| Transition | Rank | Name | Threshold | Accuracy | F1 | Progress |
| --- | --- | --- | --- | --- | --- | --- |
| ESC→EPI | 1 | Klf4 | 10.94 | 0.9 | 0.79 | $X \leq T$ |
| | 2 | Esrrb | 10.69 | 0.97 | 0.93 | $X \leq T$ |
| | 3 | Zfp42 | 13.96 | 0.93 | 0.88 | $X \leq T$ |
| | 4 | Nr5a2 | 8.99 | 0.9 | 0.79 | $X \leq T$ |
| | 5 | Tbx3 | 10.97 | 0.9 | 0.77 | $X \leq T$ |
| EPI→NPC | 1 | Gdf3 | 7.78 | 0.87 | 0.86 | $X \leq T$ |
| | 2 | Pou5f1 | 7.33 | 0.9 | 0.9 | $X \leq T$ |
| | 3 | Fgf4 | 1.9 | 0.93 | 0.92 | $X \leq T$ |
| | 4 | Dnmt3b | 16.35 | 0.88 | 0.86 | $X \leq T$ |
| | 5 | Dppa4 | 9.35 | 0.77 | 0.77 | $X \leq T$ |

For example, for the ESC→EPI transition, ReDX identifies *Klf4, Esrrb, Zfp42, Nr5a2* and *Tbx3* as the genes best able to discriminate between the cell classes; all of which have been shown to regulate naive pluripotency. After training, the learnt model identifies that thresholding for *Klf4* at 10.94 splits the cell classes such that samples whose *Klf4* expression satisfies *Klf*4 ≤ 10.94 are classified as embryonic stem cells and those for which *Klf*4 > 10.94 are classified as epiblast like cells. The directionality learnt by the model is consistent with our understanding of *Klf4*’s role in murine neuronal development, whereby it regulates key transcription factors during embryonic development and prevents the differentiation of embryonic stem cell populations [27]. *Essrb* is known to prevent the transition from ES cell to epiblast like cells through co-recruitment of *Sox2* and downstream positive regulation of NR0B1 inhibiting the epi-bast like transcriptional program [28]. This supports the features identified by RedX for characterising boundaries between cell types, and highlights the pragmatism that the simple, interpretable model structure offers for hypothesis generation.

Across both transitions, we find that the simple decision stump models exhibit high predictive performance on withheld test data. We observe a mean classification accuracy of 92% across the top 5 ranked decision stumps for the ES→EPI boundary and 87% on the EPI→NP boundary. As detailed below, section, ReDX ranks the candidate decision stumps by their prospective information gain, i.e. how effectively they segregate classes. An alternative approach would be to simply rank models using performance metrics such as the misclassification rate, evaluated on some withheld test or validation set. However, ranking by information gain alone is more intuitive and eliminates concerns with constructing a representative validation set, to which the models might otherwise overfit. As expected, we observe that high ranking models (those with high information gain) also have high predictive performance on withheld test data.

To this end, we demonstrate our method’s ability to learn simple, interpretable models from single-cell gene expression data. Further, we emphasise the utility in selecting maximally discriminant features for differentiating (in the common sense) between cell types – without the need for prior expert information e.g. feature importance or underlying noise distributions. The low-dimensional models learnt in this approach offer a framework from which we can begin to learn aspects of developmental dynamics from transcriptomic data.

### Characterising early embryogenesis in *Xenopus tropicalis*

In addition to linear developmental processes, we are also able to characterise *branching* cell fate decisions[15] using ReDX. We illustrate this in the context of early *Xenopus tropicalis* embryogenesis[29]. Here ‘Stage 08 blastula’ cells commit to one of six more specialised ‘Stage 10’ progenitors {‘Stage 10 neuroectoderm’, ‘non-neural ectoderm’, ‘marginal zone’, ‘Spemann organizer (mesoderm)’, ‘Spemann organiser (endoderm)’ and ‘endoderm’}. Applying RedX, we are able to identify the topmost informative features and thresholds for each developmental trajectory [Table 2].

**Table 2:** Repeated Decision Stumping characterises cell fate commitment of stage 08 Xenopus tropicalis blastula cells to each of the known stage 10 progenitor cell types.

| Transition | Rank | Name | Threshold | Accuracy | F1 | Progress |
| --- | --- | --- | --- | --- | --- | --- |
| SO8-blastula→S10-neuroectoderm | 1 | *Ptmap12* | 19.23 | 0.93 | 0.9 | $X > T$ |
| | 2 | *Eef1a1* | 12.12 | 0.89 | 0.83 | $X > T$ |
| | 3 | *Cirbp* | 23.37 | 0.89 | 0.83 | $X > T$ |
| | 4 | *Gsk3b* | 158.96 | 0.85 | 0.77 | $X \leq T$ |
| | 5 | *Sox2* | 2.21 | 0.84 | 0.77 | $X > T$ |
| SO8-blastula→S10-non-neural ectoderm | 1 | *Ptmap12* | 17.51 | 0.94 | 0.9 | $X > T$ |
| | 2 | *Eef1a1* | 12.06 | 0.89 | 0.83 | $X > T$ |
| | 3 | *Cirbp* | 21.39 | 0.87 | 0.79 | $X > T$ |
| | 4 | *Muc5b* | 4.69 | 0.89 | 0.85 | $X > T$ |
| | 5 | *Hnrnpa1* | 8.76 | 0.88 | 0.81 | $X > T$ |
| SO8-blastula→S10-marginal zone | 1 | *Eef1a1* | 13.42 | 0.85 | 0.79 | $X > T$ |
| | 2 | *Gsk3b* | 147.08 | 0.84 | 0.81 | $X \leq T$ |
| | 3 | *Ccnb1* | 0.69 | 0.83 | 0.73 | $X \leq T$ |
| | 4 | *Cirbp* | 21.27 | 0.82 | 0.75 | $X > T$ |
| | 5 | *Fth1b* | 1.87 | 0.76 | 0.7 | $X > T$ |
| SO8-blastula→S10-Spemann organizer (mesoderm) | 1 | *Chrd1* | 6.42 | 0.95 | 0.97 | $X > T$ |
| | 2 | *Lhx1* | 3.73 | 0.96 | 0.98 | $X > T$ |
| | 3 | *Opn7b* | 0.81 | 0.95 | 0.97 | $X > T$ |
| | 4 | *Eef1a1* | 16.39 | 0.94 | 0.96 | $X > T$ |
| | 5 | *Gsk3b* | 141.24 | 0.91 | 0.95 | $X \leq T$ |
| SO8-blastula→S10-Spemann organizer (endoderm) | 1 | *Frzb2* | 1.43 | 0.94 | 0.96 | $X > T$ |
| | 2 | *Gsk3b* | 132.51 | 0.91 | 0.94 | $X \leq T$ |
| | 3 | *Frzb1* | 2.61 | 0.88 | 0.92 | $X > T$ |
| | 4 | *Eef1a1* | 15.55 | 0.91 | 0.94 | $X > T$ |
| | 5 | *Opn7b* | 1.82 | 0.87 | 0.92 | $X > T$ |
| SO8-blastula→S10-endoderm | 1 | *Frzb2* | 3.39 | 0.97 | 0.99 | $X > T$ |
| | 2 | *Frzb1* | 8.07 | 0.96 | 0.98 | $X > T$ |
| | 3 | *Atp1a1* | 9.5 | 0.97 | 0.98 | $X > T$ |
| | 4 | *Tmem150b* | 2.45 | 0.98 | 0.99 | $X > T$ |
| | 5 | *Sox17a* | 5.61 | 0.97 | 0.98 | $X > T$ |

As with the murine results, models trained on the *Xenopus* data and ranked by information gain perform well on withheld test data. The learnt decision stumps show high mean predictive performance across 5-fold cross-validation on unseen test data, with the top models averaging 92.5% accuracy. While the performance of the highest ranked models is generally high, for some boundaries the drop-off in performance - even within the top 5 ranked models - is more pronounced. The stage 08 blastula to stage 10 marginal zone boundary, for instance, remains well characterised by EEF1AS (86% accuracy), but by rank 5, thresholding for FTH1B classifies the withheld data with only 76% accuracy. This may indicate that only a few genes robustly drive (or reflect at the level of mRNA) cell fate commitment; perhaps even more likely is that such boundaries between some cell-types are more complex than those between others. As the suggested genes are derived directly from classifiers learnt from data, we can use the model performance on unseen data to inform our belief in a gene’s involvement in a given process.

Looking for candidate developmental genes, we see that both the Neuroectoderm and Non-neural ectoderm branches returned *ptmap12* as the most informative gene (albeit with varying thresholds); this gene has long been known to be involved in *Xenopus* development [30, 31]. Thus a role in this transition is not unexpected. The overlap in discriminatory genes between the two branches, extending to EEF1AS – an elongation factor with known developmental roles in different organisms – and CIRBP – required for gastrulation, cell migration during embryogenesis, and neural developement in *Xenopus* – suggests shared, or partially shared, regulatory mechanisms for the commitment of stage 9 blastula cells to stage 10 neuroectoderm and non-neural ectoderm cell types. From previous studies[32, 33], the involvement of *gsk3b*, in several transitions is also not surprising. It appears to be required at high mRNA levels in the stage 08 blastula. and a decrease in expression is observed in several S10 stages. Because of its complex interactions[34], GSK3B, can attain multiple stable states [35], but here (see Table 2 and Figure 2), at the transcriptional level only two states can be discerned. Similarly, *frzb1, frzb2*, and *sox17a* show strong prior association with developmental processes and embryogenesis. The approach developed here yields these targets in a purely inductive, data-driven, manner without any further intervention or the requirement for domain expertise.

**Figure 1:**
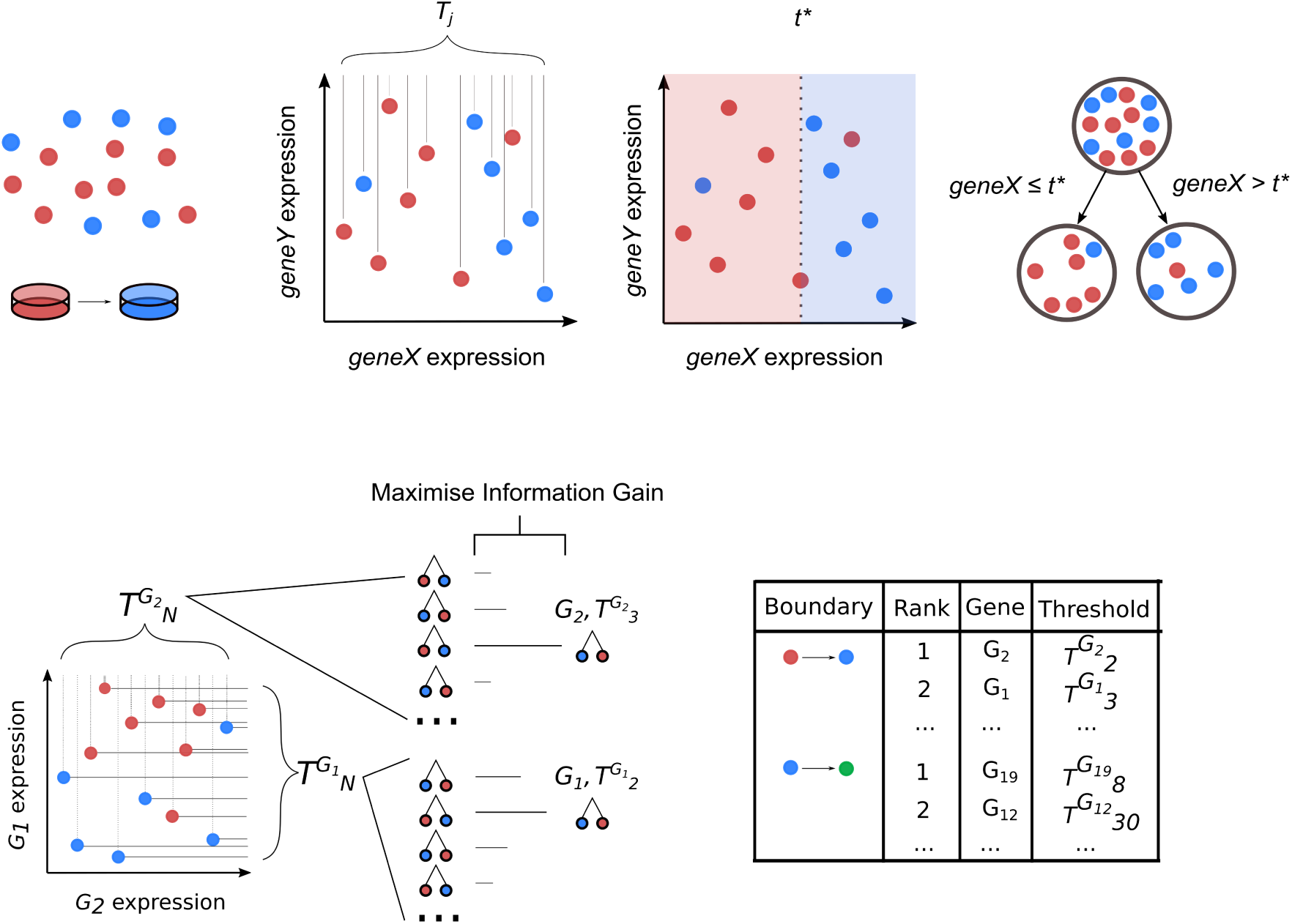
Identifying informative cell fate markers from single-cell transcriptomic data using Repeated Decision Stumping. Cells sampled across a developmental boundary are profiled and decision stump models generated to classify cell stages, using the measured expression values for each gene as potential thresholds. The resulting decision stumps are then ranked by the information gain associated with dividing the set of classes along the corresponding boundary, reporting the topmost informative genes.

**Figure 2:**
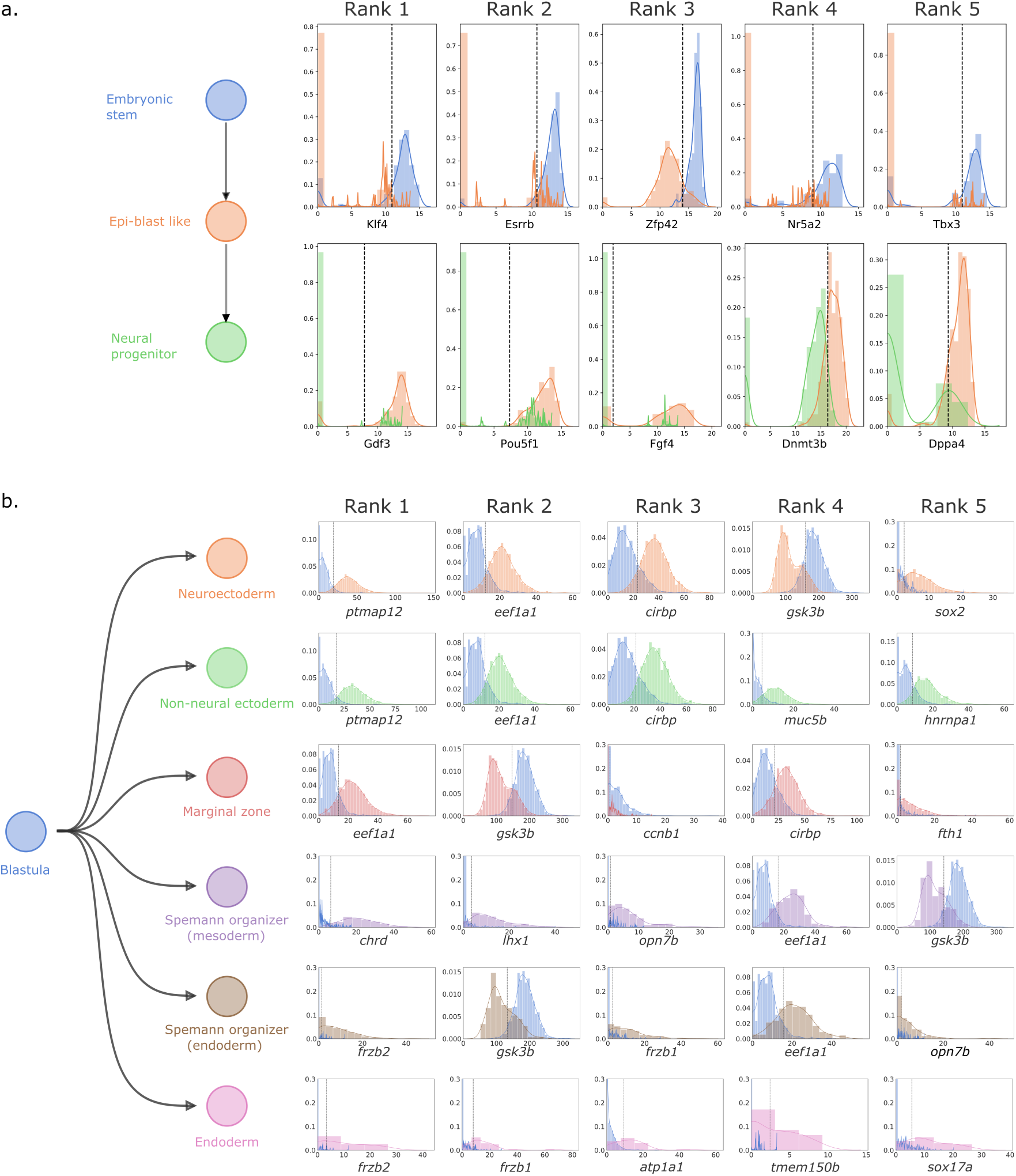
Repeated Decision Stumping identifies top features and associated thresholds for cell fate choice. a. Top five features identified for each boundary in the linear trajectory of murine neural cell fate commitment, b. Top five features identified for branching trajectories in early Xenopus tropicalis embryogenesis.

### Feature rankings are stable under random training-test partitions

We also demonstrate the robustness of predictions made by ReDX to perturbations in, or systematic problems with, the training data. To do so, we devise a stability metric[25], which describes how consistently and how highly a given feature is ranked over many random partitions of the training data (see methods).

We calculate this gamma score foreach feature over 100 random splits of the data into 80% training and 20% test partitions. Sorting the features by *γ* shows that RedX produces consistent gene rankings when trained on these different portions of the data [Figure 3]. Moreover, the feature reproduction is stable[24, 25], and approaches consistent reproduction across the random partitions. Together, these findings increase our confidence in the quality of the embedded feature selection for discerning developmental boundaries from such data.

**Figure 3:**
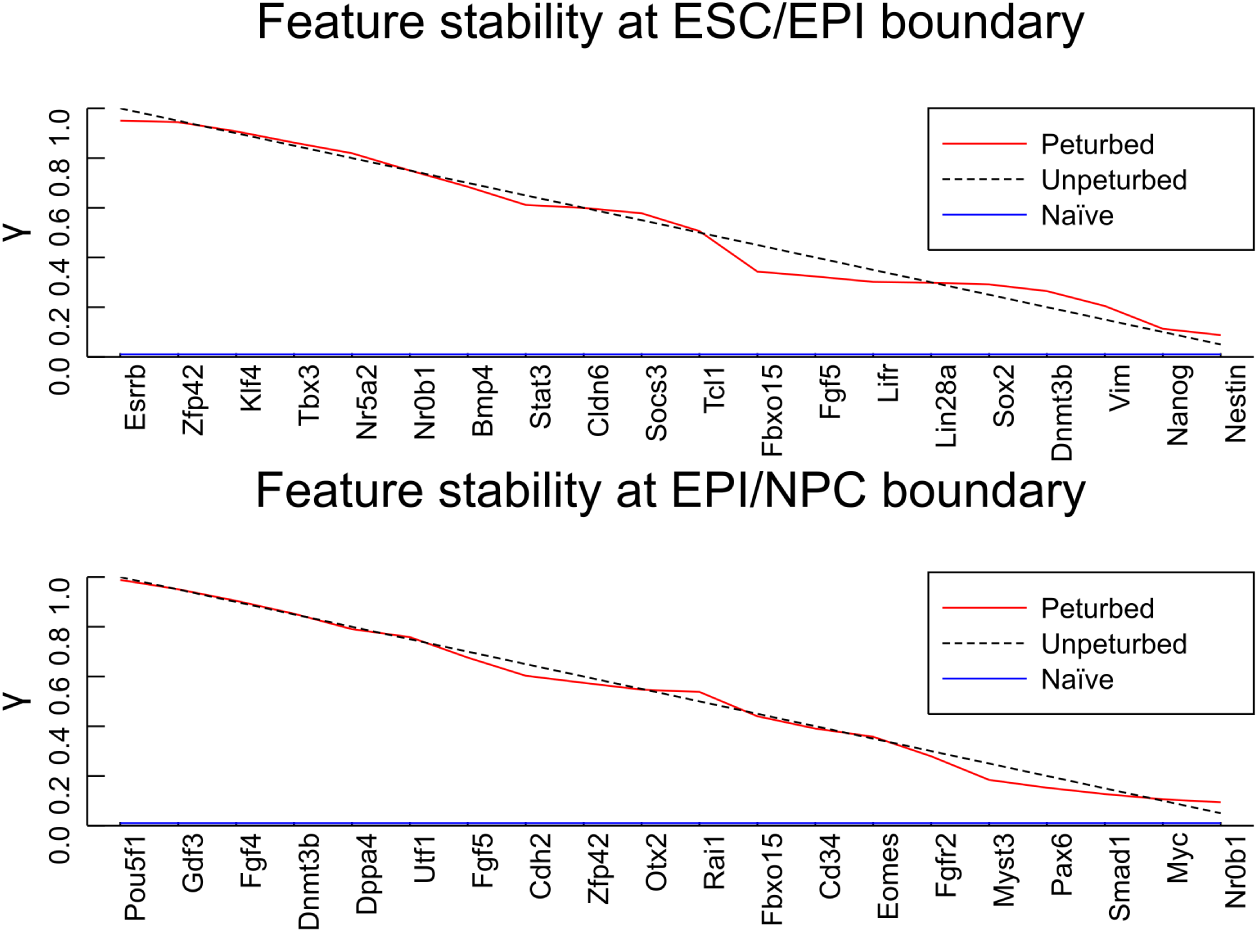
Robustness of ReDX feature rankings over 100 random partitions of the MacArthur data set into 80% training and 20% test. Features reported along the x-axis in decreasing γ. The ‘Naive’ line shows the expected γ scores if ReDX was ranking features randomly at each partition. The ‘Unpeturbed’ line shows how γ would decay if ReDX returned identical feature rankings from all 100 partitions.

**Figure 4:**
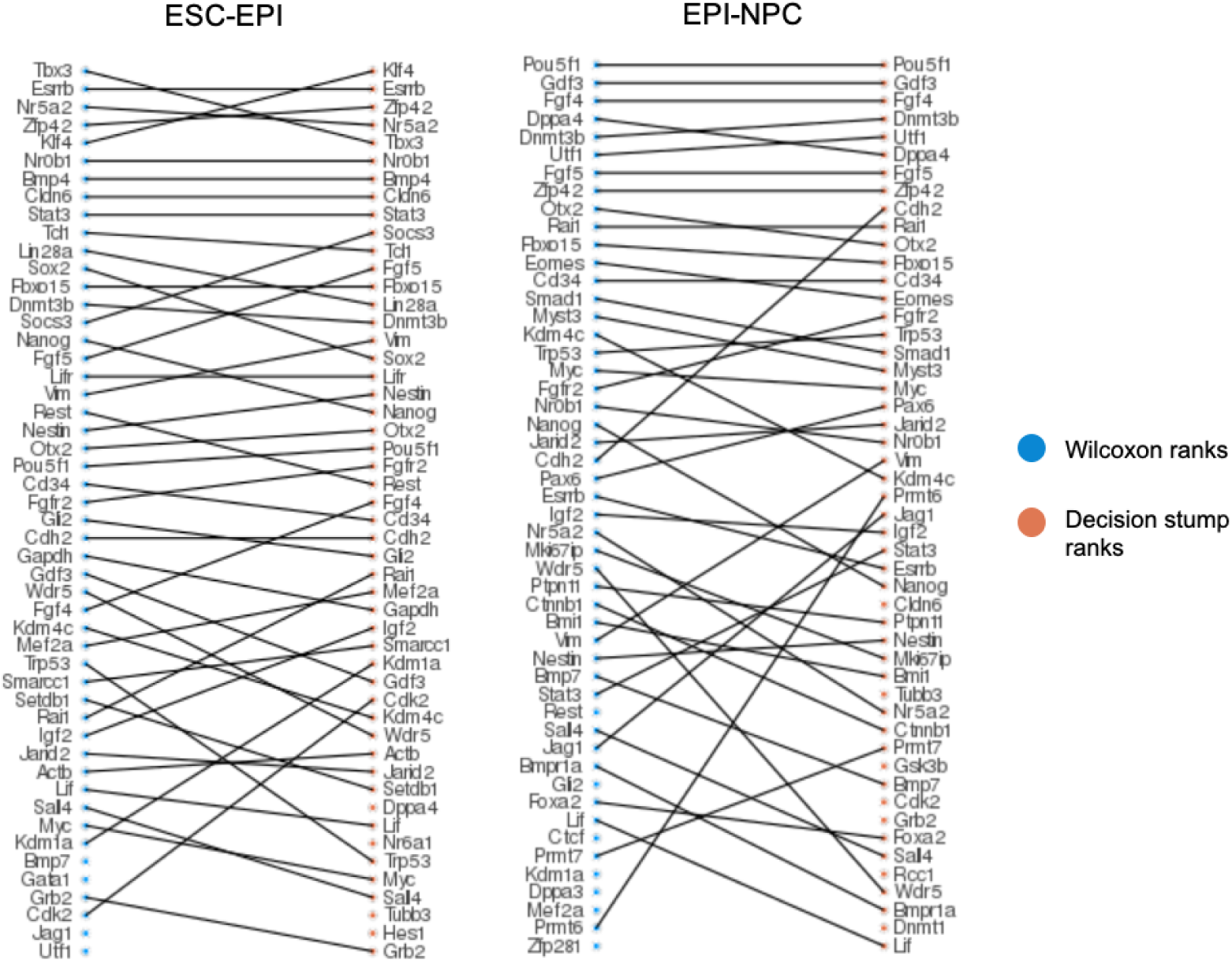
Comparison of the top 50 ranked features in the murine data following gene set enrichment analysis by Wilcoxon Rank Sum tests and Repeated Decision Stumping.

**Figure 5:**
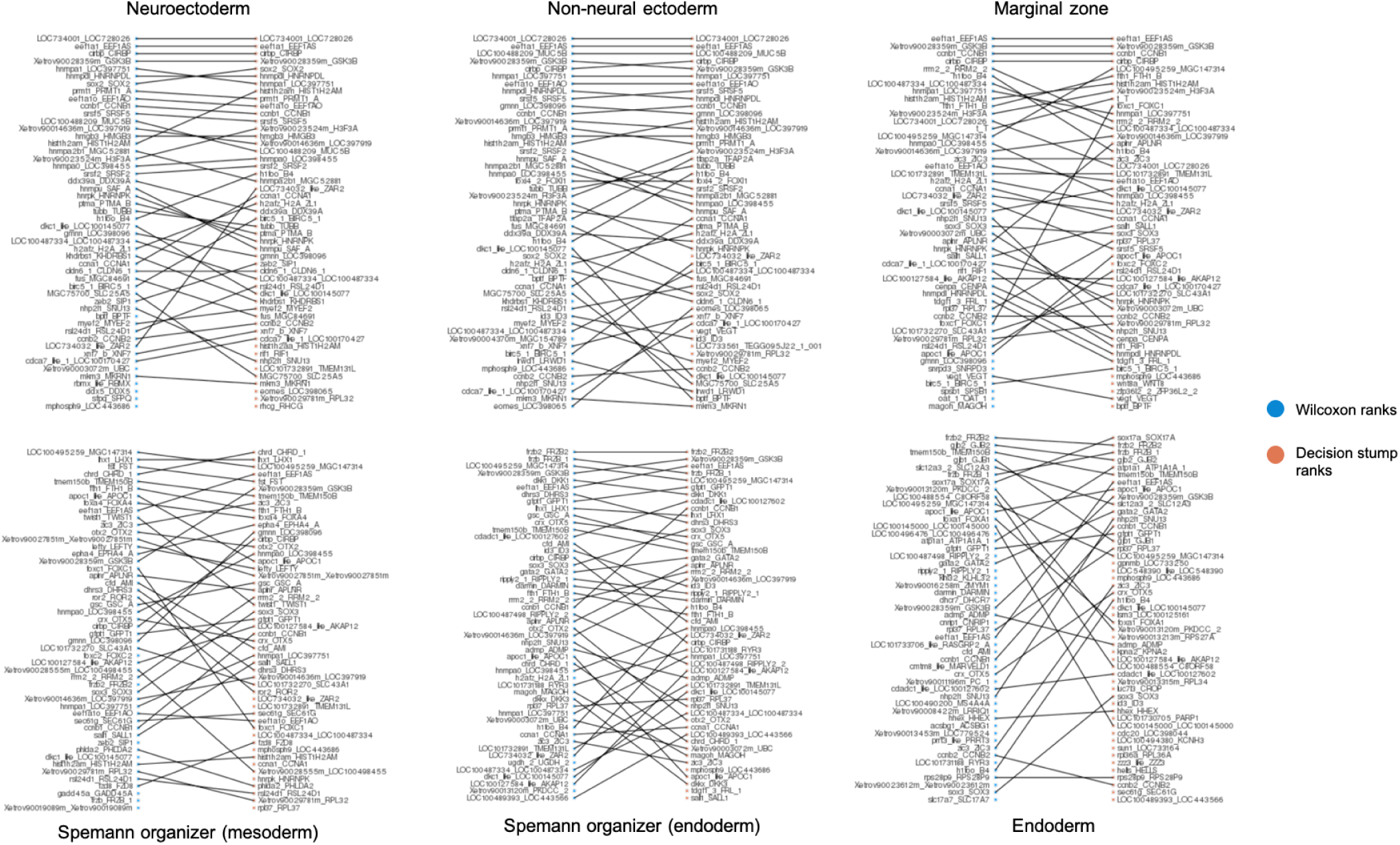
Comparisonofthetop 50 rankedfeaturesinthe Xenopustropicalisdatafollowinggenesetenrichmentanalysis by Wilcoxon Rank Sum tests and Repeated Decision Stumping.

## Conclusions

We have developed ReDX as an approach that allows us to learn *interpretable* models from high-dimensional and potentially complex data, such as current single-cell data. It is aimed to augment the impressive state-of-the-art machine learning approaches[36], such as deep neural networks and reinforcement learning, by focusing on the generation of deliberately simple mechanistic hypotheses[37, 26]. Despite the focus on informative classifiers, the features learned by ReDX also retain high predictive power. This makes them immediately applicable for informing mechanistic model development, and for generating interpretable and testable hypotheses about e.g. the underlying developmental processes. The opportunities presented by RedX are two-fold: to generate predictive models from data (which themselves can be used to annotate unlabelled samples, or identify potentially mislabelled samples); and to provide an effective, scalable means of feature selection for guiding further, (including hypothesis-driven) experimental design and mechanistic modelling.

One common critique[18] of decision trees and stumps is their instability to variations in the training data. To address this concern, we devised a quantitative stability measure, *γ*[24, 25]. Calculating it over random partitions of the training data, we are able to show that ReDX generates consistent and stable feature rankings for our developmental case studies. Whilst this may not be true of data with highly intra-heterogeneous classes, *γ* scores provide us with an objective and intuitive means of assessing the quality of predictions made using ReDX in new contexts with little available prior information.

Whilst each learnt model is simple and 1-dimensional, our findings demonstrate their ability to segregate cells along diverse developmental boundaries with great precision. Further, as a pragmatic method for gaining insights, maybe even intuition, about a high-dimensional system – all without prior expert knowledge – the merits of such an approach seem clear, especially if deployed alongside other, predictive data-driven methods.

ReDX can be applied in many contexts where labelled, single-cell data are available. Our method robustly proposes features and thresholds that optimally discriminate between the target labels (here cell states). We demonstrate the approach’s effectiveness for characterising cell-stage boundaries during healthy ontogenesis across linear and branching trajectories. The method can also applied to other, e.g. multi-omics[38, 39] without further modification (proteomic, metabolic and transcriptomic) and carries promise in biological contexts, where suitable (labelled) data and features are available but for which mechanistic models remain elusive. We can also apply ReDX to discern discriminant features in healthy *v*.*s*. diseased samples, or between control and perturbation conditions given appropriate experimental design.

The simple candidate mechanisms generated by ReDX will then serve as seeds for developing better mechanistic models.

## Methods

### Learning genes associated with developmental processes from single-cell data

Decision trees are a commonly used class of predictive statistical model capable of learning decision boundaries from labelled data[40, 41, 42, 43]. One salient characteristic of these models is their inbuilt capacity for featureselectionduringmodeltraining. Givenhigh-dimensionalinput, a low-dimensionalmodelisreturned; and they are capable of classifying samples according to a systematically reduced subset of features. We can exploit the structure learnt by these models to not only make predictions for unclassified samples, but also to better understand the boundaries between cell classes, and to gain meaningful biological insight[11, 14].

Consider a single cell, *i*, whose expression profile for *J* genes is given by the vector **x**_*i*_. Each element of the *J*-dimensional vector **x** represents the gene expression value of the *j*th gene: *x* = (*x*_1_, *x*_2_, …, *x*_*n−*1_, *x*_*n*_). For *N* such cells, consider the matrix *X*, such that *X*_*i,j*_ is similarly the expression of gene *j* measured in cell *i*. The developmental stage of each cell is given by **y**, such that *y*_*i*_ is the class label of the *i*th cell.

The first bifurcation of a decision tree splits these *N* samples into two groups, *q*_1_ and *q*_2_, using a feature, *j* and threshold *t*, such that *j* ≤ *t* for all samples in *q*_1_ and *j* > *t* for all samples in *q*_2_; these trees are referred to as *decision stumps*[17, 40]. We wish to identify the best gene and corresponding expression threshold, *j*^*∗*^ and *t*^*∗*^, respectively, with which to divide the samples such that class labels in the resulting groups are maximally segregated. If we are able to learn informative feature-threshold pairs from data, we will can gain insights into the underlying developmental dynamics of our system.

To quantify the reduction in entropy over the class labels **y** following a split at the proposed gene and threshold pair (*j, t*) we use the *information gain* (IG) measure,

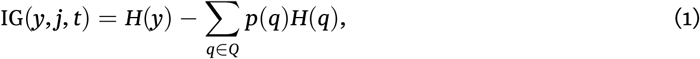

where *H*(*y*) is the entropy associated with the class labels *y, Q* is the set of subsets *q* proposed by splitting *y* on the feature *j* and threshold *t*, which thus satisfies *y* = ∪_*q*∈ *Q*_, *p*(*q*) is the fraction of elements contained in the subset *q* compared to the set *y* and *H*(*q*) is the entropy over the class labels contained in subset *q* (see methods for details).

Following established procedure for growing decision trees [43] we calculate IG over all possible thresholds for each gene (*t*_*j*_ ∈ *X*_*j*_), and select the gene and threshold for which IG is maximised (*j*^*∗*^ and *t*^*∗*^ respectively); optimising over all pairs (*j, t*) according to

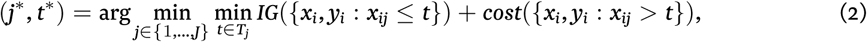

where *T*_*j*_ is the set of possible thresholds for feature *j* obtained by sorting the unique values of *X*_*j*_.

This first bifurcation of a decision tree represents the most informative axis-parallel split in the feature data for segregating the set of class labels. Intuitively, this implicit feature selection can be exploited to identify those features to which the classification problem is most sensitive.

While traditional decision tree algorithms [43], such as `ID3` and `C4.5`, recurse on the resultant leaf nodes to grow a full decision tree [44], here we interrupt the process after the first node is grown. The feature associated with the resulting *decision stump* [17] is recorded and removed from the set of available features. A new stump grown on the *full* set of samples – *excluding* the previously selected feature – thus identifies the second most informative feature/threshold pair with respect to the cell labels. This process can be repeated *n* times to identify the top *n* most discriminatory features associated with a binary classification problem. This forms the basis of our approach, for which a visual summary is provided in Figure 1.

### Learning and ranking decision stumps

To learn each decision stump, we identify the best feature, *j*^*∗*^, and best threshold for that feature, *t*^*∗*^, from labelled single-cell gene expression data. To do so, we follow the established procedure for growing a decision tree [43, 41] and optimise over all pairs (*j, t*) according to

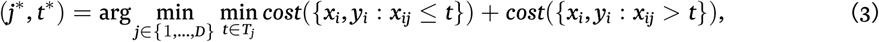

where *T*_*j*_ is the set of possible thresholds for feature *j* obtained by sorting the unique values of *x*_*ij*_. The cost function corresponds to a chosen impurity measure used to quantify the quality of a split with regards to purity of the daughter node labels. In our case, we use information gain (see below, section).

For continuous features, such as the gene expression values derived by qPCR, we consider thresholds at values for that feature already present in the training data. For each feature *j*, the cost function is calculated for all possible thresholds *t* ∈ *T*_*j*_, corresponding to the unique expression values for gene *j* across all the cell samples in the training data. As shown in equation 2, the best attribute to split on is the gene and expression threshold (*j*^*∗*^, *t*^*∗*^) that maximises label purity over the proposed daughter nodes according to the chosen cost function.

### Information Gain

Information Gain is a measure of the reduction in entropy[45, 46] in a data set given some transformation. In our case, we calculate the information gain in the set of class labels, *S*, with respect to a candidate split, *A*, according to

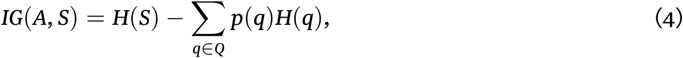

where *H*(*S*) is the entropy of the parent set *S, Q* is the set of subsets *q* proposed by splitting *S* on attribute *A* which thus satisfies *S* = ∪_*q*∈*Q*_ *q, p*(*q*) is the proportion of elements in the subset *q* compared to the set *S* and *H*(*q*) is the entropy over the subset *q*. For clarity, the entropy of any set *S* containing *C* unique classes is

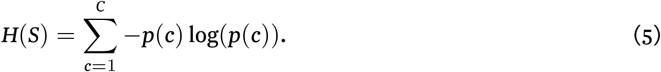

### Model evaluation

#### Misclassification rate and accuracy

After training our models, we have to evaluate the quality of their predictions on unseen data. Principally, we quantify the difference between the predicted labels and true (hidden) labels using the misclassification rate and accuracy. Let ŷ_*i*_ be the class prediction for the *i*th sample in a test set of *n* samples. Then the test set misclassification rate is simply the fraction of misclassified cases, given by,

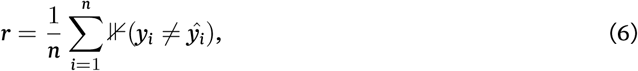

where ⊮ is the indicator function such that

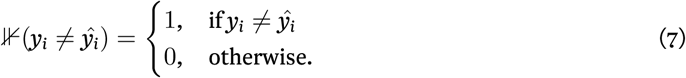

Following equation (6), the model’s accuracy on the test data is given by

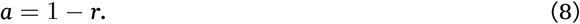

#### F_1_ score

Whilst accuracy defined in this way provides an immediately intuitive single-number evaluation metric, it fails to take into account unequal label frequencies in the test data. For instance, consider a binary classifier which simply returns label *A* every time, regardless of the input features. Using accuracy as the sole metric, we might end up considering this classifier as highly effective if the test data labels consisted of 99% label *A* and 1% label *B* (resulting in a model accuracy of 99%). However, a model which completely ignores its input features will not generalise well to new data, and should intuitively be avoided.

To address this, we also report the *F*_1_ score for each learnt model[43]. This is defined as

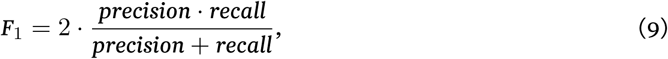

where precision is 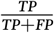 and recall is 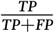.

More generally, the *F*_1_ score is a specific instance of *F*_*β*_ scores

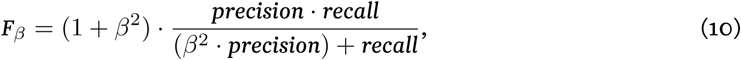

where *β* is real-valued, positive and chosen such to make recall is *β* times as important than precision. In our use cases, we have no reason to attribute more weight to either recall and precision, and report the *F*_1_ score as standard (*β* = 1).

### Feature stability analysis

One established limitation of decision trees is their instability [40]. Small variation in training data can affect the topology of the resulting tree, as well as affecting which features are selected for the discrimination of the same target variable. Decision stumps, and the order in which they are selected in ReDX, can also be affected by small changes in the training data. This is problematic if we are aiming to propose a list of the top-most informative genes to act as biomarkers, or to direct experimental and mechanistic modelling effort.

To investigate and quantify the stability of features proposed by ReDX in the context of single-cell data, we devise an associated method for calculating feature stability, following Kirk *et al*. [25], over random partitions of the training data. This stability metric, *γ*, is calculated for each feature in order to quantify how consistently genes are ranked given perturbed training data. Given a gene of interest, for *N* random partitions of the data and *q* top features for each iteration of ReDX,

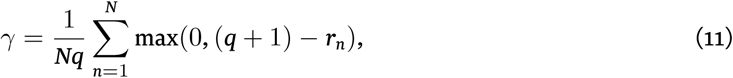

where *r*_*n*_ is the rank of that gene at the n^th^ partition.

As such, *γ* is confined to {*γ* ∈ ℝ : 0 ≤ *γ* ≤ 1}. A value of 1 would mean that the gene was selected as the most informative over every random partition of the data and a value of 0 indicates that the gene was never selected to be in the top q stumps over the random partitions. A value of 0.5 might indicate that the feature was selected in half of the partitions at rank 1 and unranked in the other half, or the feature may have been chosen for the middle rank in all of the random partitions, or some equivalent thereof.

### Datasets and preprocessing

#### Linear Differentiation Pathway

The first data set contains single-cell gene expression data for mouse embryonic stem cells undergoing differentiation to neural progenitor cells [26]. In the initial study, the mouse embryonic stem cells in the E14tg2a and R1 embryonic mouse cell lines were profiled during cell fate commitment, identifying a previously unknown epiblast like, intermediary cell state.

In total, 547 cells were sampled sampled over the course of seven days; at 24h, 48h, 72h, 120h and 168h after transfer from leukemia inhibitory factor (LIF) + 2i conditions to N2B27 neural basal medium; following existing protocol [47]. Gene expression was reported for 96 genes affiliated with ES cell development using a high-throughput RT-PCR array. The resulting 547×96 feature matrix was supplemented by additional annotations provided in the initial work for each cell sampled. These included the sampling time for each cell, alongside a classification label for the cell; “ESC”, “EPI” or “NPC”; representing populations of embryonic stem cells, epiblast-like intermediate cells and neural progenitor cells respectively. As discussed in the supplementary material of the original work, these cell class labels were assigned through k-means clustering. Together with the gene expression data, these annotations provide us with a gold standard reference with which to train the models in our own work and to evaluate their predictive performance.

#### Branching Differentiation Pathway

The second data is derived from a larger high-throughput sequencing effort, focused on early embryo development in *Xenopus tropicalis* [29]. A total of 37,136 cells were sampled from zygotic genome activation (stage 8, 5 hours postfertilization) through to early organogenesis (stage 22, 22 hours postfertilization). Single cells were profiled with sc-RNAseq, following a previously established high-throughput single-cell droplet barcoding pipeline [48]. The resulting data describes the underlying scRNA-seq counts for 25,087 genes in each of the sampled cells. Cell stage annotations were derived in the original work, using hierarchical clustering on the scRNA-seq data and cluster specific genes used to draw cluster labels from known embryonic cell types documented in the Xenopus Bioinformatics database – Xenbase [49].

The full scRNA-seq count data and raw FASTQ files remain publicly accessible at tinyurl.com/scXen2018.

## Availability of data and materials

The datasets analysed during the current study and the associated code are available in the RepeatedDecisionStumping GitHub repository, https://doi.org/10.5281/zenodo.4017262

## Author’s contributions

IACV and MPHS conceived the methodology; IACV developed the algorithm and performed the analysis; IACV and MPHS wrote the paper.

## Acknowledgements

We gratefully acknowledge the support from the members of the *Theoretical Systems Biology Group* at Imperial College London and the University of Melbourne. IACV is funded through a Wellcome Trust PhD Studentship. MPHS gratefully acknowledges funding from the Volkswagen Foundation and the Deputy Vice Chancellor of Research Fund.

## Supplementary Material

### Concordance with Wilcoxon Rank Sum tests

The Wilcoxon Rank Sum is a commonly used method in gene set enrichmentanalysis. Similarly toourmethod of repeated stumping, it is used to identify differentially expressed genes between groups of cells. It also shares its salient non-parametric form and calculations based on ordinal transformations with our decision stump approach, which combine to make the methods particularly amenable for gene expression data with potentially many outliers and whose underlying distributions are unknown.

To compare how similarly the two approaches rank enriched features, we also performed a Wilcoxon Rank Sum test for each gene across each of the developmental boundaries considered in the main body of this work. Ranking these genes by the significance level, and applying a Benjamini-Hochberg correction for multiple testing[42] using a false discovery rate of 0.01, we were then able to compare these lists of ranked surviving feature with those produced by the decision stumps.

The concordance between the established Wilcoxon Rank Sum approach and our decision stump implementation improves our confidence in the validity and potential of decision stumps for identifying gatekeeper genes. This is especially true for the *Xenopus* data, where both approaches select a highly similar set of the 50 topmost signifcant genes from 25,087 possible features. Further, we believe that the mechanistic interpretation offered by the learnt decision stump models – in terms of their learnt expression threshold/feature pair – brings additional value to initial gene set enrichment analyses than the conventional statistical enrichment methods. That the learnt models can also behave as classifiers in their own right is also of significant interest and utility e.g. labelling unclassified samples, generating binary networks.

